# A causal role for pulvinar in coordinating task independent cortico-cortical interactions

**DOI:** 10.1101/2020.03.07.982215

**Authors:** Manoj K. Eradath, Mark A. Pinsk, Sabine Kastner

## Abstract

Pulvinar is the largest nucleus in the primate thalamus and has topographically organized connections with multiple cortical areas, thereby forming extensive cortico-pulvino-cortical input-output loops. Neurophysiological studies have provided evidence for a role of these transthalamic pathways in regulating information transmission between cortical areas. However, a causal role of pulvinar in regulating cortico-cortical interactions has not yet been demonstrated. In particular, it is not known whether pulvinar’s influences on cortical networks are task-dependent or reflect more basic large-scale network properties that maintain functional connectivity across a network regardless of active task demands. In the current study, under a passive viewing condition, we conducted simultaneous electrophysiological recordings from interconnected ventral (area V4) and dorsal (LIP) nodes of the macaque visual system while reversibly inactivating the dorsal part of lateral pulvinar (dPL), which shares common anatomical connectivity with V4 and LIP. Our goal was to probe a causal role of pulvinar in regulating cortico-cortical interactions in the absence of any active task demands. Our results show a significant reduction in local field potential phase coherence between LIP and V4 in low frequencies (4-15 Hz) following muscimol -a potent GABA_A_ agonist -injection into dPL. At the local level, no significant changes in firing rates or LFP power were observed in LIP or in V4 following dPL inactivation. These results indicate a causal role for pulvinar in synchronizing neural activity between interconnected cortical nodes of a large-scale network, even in the absence of an active task state.

**Significance Statement:** Pulvinar, the largest nucleus of the primate thalamus, has been implicated in several cognitive functions. The extensive cortico-pulvino-cortical loops formed by pulvinar are suggested to be regulating information transmission between interconnected cortical areas. However, a causal evidence for pulvinar’s role in cortico-cortical interactions in the absence of active task demands is not yet clear. We conducted simultaneous recordings from nodes of macaque visual system (areas V4 and LIP) while inactivating the dorsal part of the lateral pulvinar (dPL) under a passive viewing condition. Our results show a significant reduction in local field phase coherence between LIP and V4 in low frequencies (4-15 Hz) following inactivation of dPL, thus providing evidence for a causal role of pulvinar in regulating cortico-cortical interactions even in the absence of an active task state.

## Introduction

Pulvinar is the largest nucleus of the primate thalamus and primarily part of the visual system. It has been implicated in several cognitive functions including orienting to stimuli, filtering of distracter information, and visually guiding motor actions (Robinson, 1992, Adams, 2000; Shipp, 2003; Kaas, 2007; Wilke, 2010). For example, in humans, deficits in filtering distracter information have been observed in patients with pulvinar lesions (Rafal, 1987; Ward, 2002; Arend, 2008; Snow, 2009), similar to the filtering deficits observed after posterior parietal cortex (PPC) lesions (Posner, 1984; Friedman-Hill, 2003; Han, 2004). In accordance with a role of pulvinar in attentional processing, monkey physiology studies have shown enhanced responses in subsets of pulvinar neurons following spatial cues that direct attention to a location in the visual field in covert attention tasks (Petersen, 1987; Robinson, 1992; Saalmann, 2012; Zhou et al.; Fiebelkorn, 2019a) or in perceptual suppression tasks (Wilke, 2009). Deficits in shifting attention to the contra-lesional visual field (Petersen, 1987; Desimone, 1990) and in performing visually guided movements into the contra-lesional space have been observed as a consequence of pulvinar inactivation (Wilke, 2010). Thus, the results from lesion and electrophysiology studies indicate an important role of pulvinar in attentional visuo-spatial processing.

Directly connected cortical areas are generally interconnected via pulvinar through topographically organized, layer-specific feedforward-feedback connections, forming an extensive network of cortico-thalamo-cortical pathways (Shipp, 2003). Pulvinar subdivisions have been shown to reciprocally connect with the frontal eye field (FEF) (Trojanowski, 1974; Gutierrez, 2000), cingulate and retrosplenial cortex (Watson, 1973; Baleydier, 1985), PPC and extrastriate visual cortex (Yeterian, 1997; Adams, 2000; Shipp, 2001; Shipp, 2003; Gattas, 2014; Arcaro, 2015b). It has been suggested that these cortico-thalamo-cortical loops form indirect (transthalamic) pathways through which the pulvinar may influence cortical processing (Kastner, 2000; Gutierrez, 2000; Adams, 2000; Shipp, 2001; Shipp, 2003; Gattas, 2014; Arcaro, 2015b; Jaramillo, 2016). In line with the cortico-thalamo-cortical loop hypothesis, electrophysiological studies have shown that the transthalamic pathways serve to regulate information transmission between interconnected cortical areas (Saalmann, 2012; Halassa, 2017; Fiebelkorn, 2019a). Specifically, during selective visual attention, the pulvinar synchronizes neural activity between interconnected cortical areas in the alpha/low beta frequency range (8-15Hz), as shown for visual areas V4 and TEO (Saalmann, 2012), the lateral intra parietal area (LIP) and V4 (Saalmann, Ly, 2018) and LIP and FEF (Fiebelkorn, 2019a, b). However, it is not clear from these correlational studies whether (i) the pulvinar *causally* influences cortico-cortical interactions and (ii) whether these interactions depend on active task demands.

While inactivation studies of the pulvinar have supported the notion of a causal role of pulvinar in spatial attention and visually guided motor actions by demonstrating behavioral deficits (Petersen, 1987; Desimone, 1990; Wilke, 2010), only few studies have examined the causal impact of pulvinar inactivation on cortical circuitry by simultaneously recording from cortical areas while selectively altering neural activity in the pulvinar. In anesthetized preparations, it has been shown that focal pharmacological inactivation of the lateral pulvinar significantly reduced evoked visual responses in V1, while focal pharmacological excitation significantly enhanced them (Purushothaman, 2012). In awake, behaving monkeys, muscimol inactivation of the ventral portion of the lateral pulvinar caused reduction in attention-related effects in V4 such as decreases in visually-evoked responses and gamma synchrony (Zhou, 2016), and, possibly as a consequence of these local effects, reduced functional connectivity between V4 and IT cortex (Zhou, 2016). Both the behavioral and electrophysiological effects following pulvinar inactivation suggest a causal influence of pulvinar on interconnected cortical areas (Zhou, 2016; Halassa, 2017). However, it is not clear whether the pulvinar influences on cortical areas were active task-related or reflected more basic properties of a large-scale network that maintains functional connectivity regardless of specific task state. Here, we asked whether the pulvinar plays a broader role in regulating functional connectivity across cortical networks by examining the pulvinar’s causal influences on interconnected cortical areas under a passive viewing condition. We conducted simultaneous recordings from nodes of the visual system (areas V4 and LIP) while inactivating the dorsal part of the lateral pulvinar (dPL), which shares common connectivity with V4 and LIP, while monkeys passively viewed videos on monitor, to explore a causal role of pulvinar in influencing cortical activity and cortico-cortical interactions in the absence of an active task structure.

## Materials and Methods

### Animals

Two male macaque monkeys (*Macaca fascicularis*; monkey B and monkey R, age 8 and 14 years, respectively) were used for the study. Princeton University Animal Care and Use Committee approved all surgical and experimental procedures, which conformed with the National Institute of Health guidelines for the humane care and use of laboratory animals.

### Identifying and accessing the interconnected target areas

We performed simultaneous electrophysiological recordings and pharmacological interventions from anatomically and functionally interconnected distant sites. The macaque D99 digital template brain atlas (Reveley, 2016) was non-linearly warped to the individual animal’s structural MRI images to outline the anatomical locations of the cortical areas of interest and pulvinar subdivisions in the individual animal’s anatomical space. The structural connectivity between pulvinar subdivisions and the cortical target areas V4 and LIP was confirmed in one animal with diffusion MR imaging (DMRI). Data were collected for the whole brain on a Siemens 3T MAGNETOM Prisma (80 mT/m @ 200 T/m/s gradient strength) using a 4-channel flexible coil or a surface coil (4-ch Flex Coil Small; 11cm Loop Coil; Siemens AG, Erlangen Germany). Diffusion images were acquired using a double spin-echo EPI readout pulse sequence with the 4-channel coil wrapped around the head. Two datasets of 270 gradient directions were collected from monkey B using a monopolar gradient diffusion scheme with *b* values distributed optimally across 3 spherical shells (field of view (FOV) = 96mm, matrix size (MS) = 96×96, slice thickness = 1.0mm, in-plane resolution = 1.0 × 1.0mm, slice orientation/order = coronal/interleaved, phase-encode directions = RL LR, number of slices = 50, repetition time (TR) = 6800ms, echo time (TE) = 66ms, phase partial Fourier = 6/8, iPAT = 2 (GRAPPA), fat suppression = on, *b*-values = 850, 1650, 2500 s/mm^3, gradient directions per shell = 90, acquisition time (TA) = 31 min 44 sec). Eighteen non-diffusion weighted images were collected interspersed throughout each dataset. After DMRI pre-processing (de-noising (Veraart, 2016; Tournier et al. 2019), Gibbs ringing artifact removal (Perrone et al. 2015; Kellner et al. 2016), susceptibility-, eddy current-, and motion-induced distortion corrections (Andersson et al. 2003), probability distributions of fiber direction at each voxel were derived with FSL (FMRIB, Oxford, UK)’s “Bedpostx GPU” tool (Hernández, 2013) considering three fiber populations per voxel (Jbabdi,2012; Behrens, 2003; Behrens, 2007). These distributions, which take into account fiber direction uncertainty due to MR noise, artifacts, and incomplete modeling of the diffusion data, were used to perform pair-wise mask tractography in order to determine likely fiber pathways between the pulvinar and each cortical area (LIP, V4). Only tracts that passed through the internal capsule were retained. In addition, tracts that entered the opposing hemisphere, the optic nerves, or the external capsule, as well as tracts that extended anterior to the pulvinar were discarded with exclusion masks. Once probable fiber pathways between the pairs were defined, they were thresholded to remove the lower 1% of probable tracts and masked by the pulvinar ROI to delineate cortical projection zones in the pulvinar. We targeted the portion of the lateral pulvinar that showed shared projection zones with LIP and V4 (**Fig. 1-1**). Averaged high resolution structural scans were used to determine the locations of recording chamber placement and sites for the craniotomies in order to access the brain sites of interest (dorsal portion of lateral pulvinar (dPL), LIP and area V4). Within the chambers, craniotomies of ∼4.5 mm size were performed above the areas of interest, while leaving the dura intact. A custom prepared MR-compatible grid tube consisting of a bundle of polyimide tubes was then inserted into each craniotomy to seal the holes, while allowing permanent access to the recording sites. The top of the grid tubes (12-20mm in total length) was then sealed with removable bio-compatible silicon adhesives to maintain the aseptic environment. We inserted tungsten micro-electrodes into the cortical areas (LIP and V4) and performed structural MRI scans to verify the electrode locations (**Fig. 1-2**) by aligning the images with D99 atlas (Saleem and Logethetis, 2007; Reveley, 2016).

### Pharmacological inactivation and visualization of the inactivation zone

Muscimol is a γ-aminobutyric acid (GABA_A_) receptor agonist and its potent central nervous system (CNS) depressant property has been used to reversibly inactivate brain regions of interest. Muscimol rapidly induces a local hyperpolarization around the injected site (Petersen, 1987; Majchrzak and Scala, 2000). A custom prepared injectrode assembly (Plastics One Inc, Roanoke, VA) was used to deliver muscimol and to simultaneously record the signals from the lateral pulvinar. The injectrode assembly consisted of 27G fused Silica cannula (od: ∼410 μm) with a central core of an epoxy coated tungsten electrode (∼125 μm). The cannula was positioned based on the measurements from the structural and diffusion MR images. We used Gadolinium, a MRI contrast agent (Magnevist, 5mM solution), to visualize (i) the initial injection site and (ii) approximate the time course of the spreading of the agent around the initial site of injection. At the beginning of each session, the monkey was sedated with a combination of ketamine (5mg/Kg) and xylazine (1mg/Kg), and the injectrode was inserted into the dPL through the grid tube. An MR scan was acquired to confirm the initial position of the injectrode by visualizing its tip. Then, 0.5 μl of Gadolinium was pressure delivered through the injectrode using a gas-tight Hamilton syringe (10μl capacity) at a rate of 0.05μl/min with the help of an automated drug delivery pump (Harvard Apparatus, MA). The macaque D99 digital template brain atlas was non-linearly warped to the animal’s MR images to confirm the initial site of the Gadolinium contrast agent injection within the dorsal portion of the lateral pulvinar (**Fig. 1A**). After the animals recovered from the sedation, they were taken to the experimental setup for electrophysiological recordings. 1μl of muscimol solution in sterile Phosphate buffered Saline (PBS) with an effective muscimol concentration of 6.67μg/μl or 1μl of sterile PBS was pressure delivered through the injectrode during the inactivation and control sessions, respectively. We performed 12 muscimol inactivation sessions (8 in monkey B and 4 in monkey R) and 10 saline control sessions (7 in monkey B and 3 in monkey R). In monkey B, we also tracked the spread of the contrast agent from the initial point of contrast injection with the help of multiple repeated structural scans over a period of 3 hours after an injection of 1.0μl of Gadolinium contrast agent into dPL. 10 T1-weighted volumes were collected (at times 0, 15, 30, 45, 60, 75,90,105,120,and 180 minutes from the end of injection), with the surface coil secured above the head to obtain high-quality structural volumes (3D magnetized prepared rapid gradient echo (MPRAGE) sequence; FOV = 128mm, MS = 256×256, slice thickness = 0.5mm, in-plane resolution = 0.5 × 0.5mm, slice orientation/order = sagittal/interleaved, TR = 2700ms, TE = 2.32ms, inversion time = 850ms, flip angle = 9 deg, number of slices = 240, TA = 11 min 21 sec). These data provided an approximation of the spread of injected substance over the course of a recording session (**Fig. 1B**). Based on the data of the temporal spread of the Gadolinium contrast agent within the pulvinar, two windows of interests were identified. A 15-minute window immediately before the start of injection (muscimol or PBS control) was used as pre-injection baseline window while a 15-minute window starting from 15 minutes after the end of injection (indicated by orange bar in **Fig. 1B**) as the post-injection window. The spread of gadolinium contrast agent was approximately within the anatomical boundaries of dPL (a volume 3mm3) during this 15-minute post-injection window (15-30 minutes from the end of injection).

**Figure 1.**
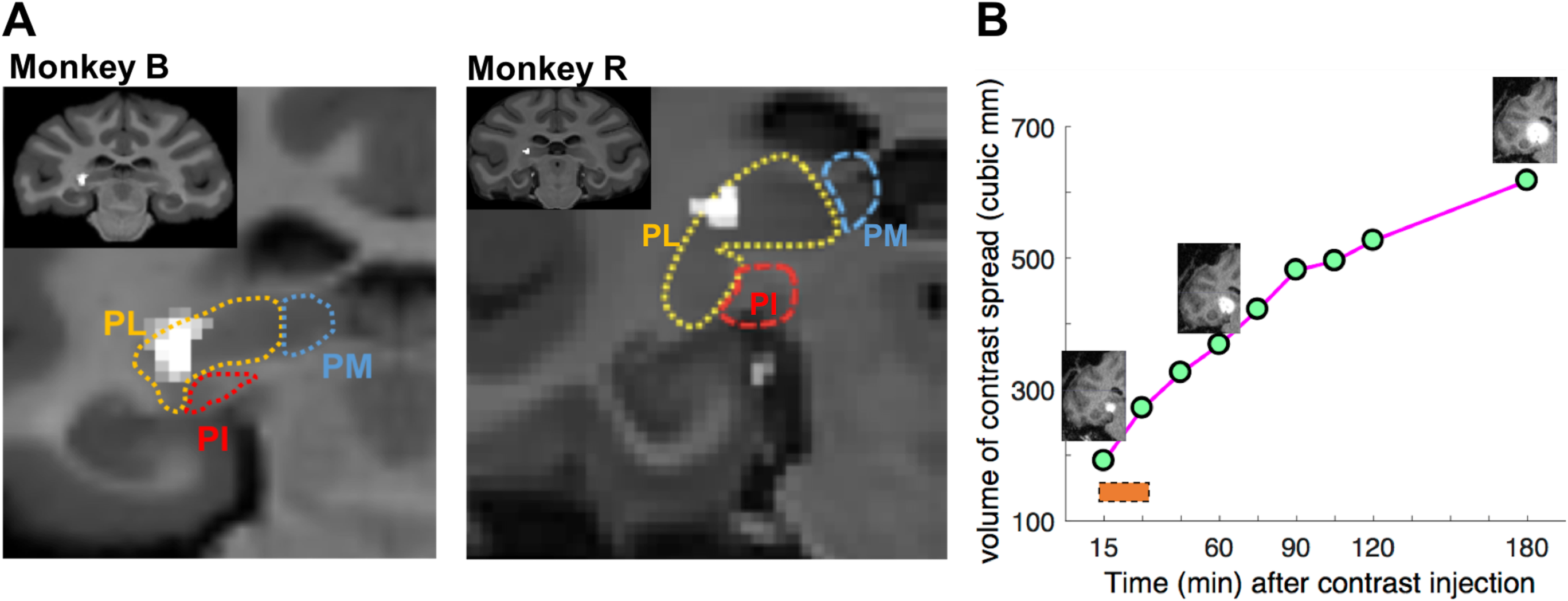
Anatomical localization of inactivation zone. (A) Structural MRI scan visualizing the initial site of a Gadolinium MRI contrast agent injection (0.5μl) in monkey B and monkey R. MR images of individual animals were non-linearly wrapped onto the D99 digital template atlas to approximate the boundaries of pulvinar subdivisions. PM: Medial pulvinar, PL: Lateral pulvinar, PI: Inferior pulvinar. The injection site was localized within dorsal portions of the lateral pulvinar (dPL) for both monkeys. (B) Serial MRI images showing the spread of Gadolinium contrast agent over a duration of 3 hours, following injection into dPL. The inserts visualize the spread of the contrast agent at 15, 60, and 180 minutes, respectively, after the end of the contrast injection, during one session in monkey B. The orange bar indicates the time window (15-minutes, starting from 15 minutes after the end of injection) used for the post-injection analyses of recordings.

### Electrophysiology

Epoxy coated single unit tungsten microelectrodes (FHC Inc.) were used in monkey B, while 32 channel multi-contact probes (V-probe, Plexon Inc., Dallas, TX) were used in monkey R for recordings of spiking activity and local field potentials (LFP). The electrodes were lowered at different angles and locations to each of the cortical areas using independent microdrives (NAN instruments). Electrophysiology signals were collected at a sampling rate of 40,000Hz for spikes and 1,000 Hz for LFPs, using a head-stage amplifier and recording system (Plexon Inc., Dallas, TX). A skull screw inserted over the opposite hemisphere (over primary motor cortex, AP +17, distance in mm from ear bar zero) and outside of the recording chambers was used as a reference for electrophysiological recordings. Inactivation/electrophysiological sessions were performed only after a minimum of 2 hours following the last ketamine/xylazine injection, after animals completely recovered from sedation. At the beginning of the session, the animals performed a receptive field (RF) mapping task to position the electrodes, so that RFs matched across the three areas. During the RF mapping task, visual stimuli (bright discs of 2 degrees’ visual angle in size) were sequentially shown in different quadrants of the monitor, while the animals maintained fixation at the center of the screen. The neuronal responses evoked by the stimuli were then analyzed online (Neuroexplorer, Plexon Inc., Dallas, TX) to identify the RFs of the recorded locations. An infrared eye tracker (EyeLink, 1000 Plus, SR Research Ltd, Ottawa, CAN) was used to monitor the animal’s eye positions during the RF mapping task. Recording sites were adjusted to align RF locations across all 3 recorded areas (dPL, LIP and V4). Spiking activity and LFPs were recorded simultaneously from dPL, LIP and V4 in monkey B and from LIP and V4 in monkey R. During the inactivation and electrophysiological recordings, the animals remained in a primate chair, head fixed, while freely viewing a video (short clips of documentary ‘snow monkeys’, PBS nature documentary), presented on the LCD monitor, placed in front of the chair. The same video clips were played during all recording sessions and for both animals. In order to assess any visual field deficits as a consequence of pulvinar inactivation/injection, monkeys performed a visually guided saccade task at the end of the session. In this task, the animals directed their gaze to and maintained fixation for 500ms at spatial cues, presented sequentially in different locations of the visual field. The animals did not show visual field deficits following inactivation/injection of dPL according to this test. Simultaneous LIP and V4 recordings were performed in both animals, while additional simultaneous pulvinar recordings during injection sessions were obtained only from monkey B (n=8, muscimol sessions, n=7, saline control sessions).

### Data analysis

#### LFP power and spike rates

Powerline noise was removed from the LFP signals using a Butterworth notch filter (filter designed using the *designfilt* function of MATLAB with notch defined as the 59 to 61 Hz frequency interval). The Hilbert transform was then applied to the LFP data to obtain instantaneous power over a frequency range of 2-90 Hz. The percentage change in LFP power was calculated by comparing the15 minutes post-injection window with that of the 15-minutes pre-injection window. The percentage changes between muscimol and control sessions were statistically compared using the Wilcoxon rank-sum test. Spiking activity was processed offline (Offline sorter, Plexon, Inc., Dallas, TX) to identify spike waveforms that crossed the threshold over 4 standard deviations from the mean of peak heights in the histogram. The single-unit/multi-unit (SUA/MUA) clusters were isolated by running a valley seeking algorithm, which assigned the waveforms into optimal number of clusters based on the inter-point distances in the feature space (Offline sorter, Plexon, Inc., Dallas, TX). A total of 56 LIP cells (monkey B: 20 cells, monkey R: 36 cells) and 50 V4 cells (monkey B: 10 cells, monkey R: 40 cells) were recorded during the muscimol inactivation sessions. 37 LIP cells (monkey B: 10 cells, monkey R: 27 cells) and 31 V4 cells (monkey B: 8 cells, monkey R: 23 cells) were recorded during the saline control sessions. Additionally, in monkey B, pulvinar neurons were recorded during the sessions (8 cells in muscimol sessions and 7 cells in control sessions).

#### LFP-LFP coherence analysis

Phase locking value (PLV) is a metric used for quantification of frequency-specific synchronization between LFP signals recorded from different brain areas (Lachaux, 1999) and is assumed to be a measure of inter-areal functional connectivity. PLV uses only the phase values to calculate the coherence between two simultaneously recorded signals. The Hilbert transform was used to determine instantaneous phase of LFP data binned into 1-second time segments over a frequency range of 2-60 Hz. Instantaneous phase was then used to calculate PLV, as defined by the equation 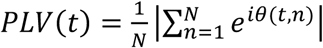, where *N* is the total number of time bins of LFP data and *θ*(*t, n*) is the difference between the instantaneous phase of two signals at time *t* and time bin *n*, in a given frequency band. The resulting PLV for each frequency was averaged over 1-second time segments to obtain frequency specific interareal PLV. The Wilcoxon sign-rank test was used to statistically compare the PLV obtained pre- and post-injection, and in muscimol and control sessions.

#### Spike-field coherence analysis

While PLV provides a measure of inter-areal synchronization, it does not inform about the directionality of the interaction. Spike-field coherence (SFC) measures the clustering of spiking activity in one area relative to specific phases of simultaneously recorded frequency-specific LFP signals from a second area, and thereby provides an alternative estimate of network connectivity between regions (Fries, 2001). Since spiking activity is generally considered to be an area’s output signal and LFP activity an area’s input signal, SFC provides indirect evidence for the directionality of inter-areal interactions (Pesaran, 2010). SFC between pulvinar and cortical areas was calculated over a frequency range of 2-60Hz, from LFP and spike data binned into 1-second time segments (*coherencycpt* script, Chronux toolbox, Chronux.org). Changes in SFC between pulvinar and cortical areas during pre- and post-injection were calculated in both directions (i.e. pulvinar spike to LIP phase, LIP spike to pulvinar phase, pulvinar spike to V4 phase, and V4 spike to pulvinar phase). The Wilcoxon rank-sum test was used to statistically compare the changes in SFC values calculated separately for muscimol and control sessions.

## Results

We performed simultaneous recordings of spiking activity and LFPs from areas LIP and V4 from awake, head-fixed monkeys while monkeys freely viewed short video clips presented on a LCD screen placed in front of the primate chair (see methods for details), before and after inactivating the dorsolateral pulvinar (dPL) by injecting the GABA_A_ agonist muscimol, or by injecting saline as a control. Based on previous studies that showed robust functional connectivity of interconnected large-scale networks even in the absence of active task requirements (Wang, 2012; Arcaro, 2015a), we used simultaneous electrophysiological recordings under passive viewing condition to probe the causal role of pulvinar in cortical networks.

### Local responses in cortical areas and pulvinar following muscimol inactivation

First, we examined local effects of muscimol inactivation on firing rates and LFP power in cortical areas LIP and V4. Firing rates in LIP did not significantly change following muscimol (pre-injection: 11.51 ± 1.15 Hz; post-injection: 11.94 ± 1.17 Hz, mean ± s.e.m., n=56; p=0.70, Wilcoxon sign-rank test) or control (pre-injection: 9.39 ± 0.67 Hz; post-injection: 10.77 ± 0.66 Hz, mean ± s.e.m., n=37; p=0.13, Wilcoxon sign-rank test) injections (**Fig. 2A, left**). LFP power in LIP before and after the injections was also not significantly different between muscimol and saline control sessions in any of the frequency ranges that were examined (theta (4-7 Hz) p= 0.97,; alpha/low-beta (7-15 Hz) p= 0.53; beta (15-30 Hz) p=0.49; low-gamma (30-60 Hz) p=0.28; or high-gamma (60-90 Hz) p= 0.93, Wilcoxon rank-sum test) (**Fig. 2A**,**right)**. Similarly, V4 firing rates did not significantly change following muscimol (pre-injection: 24.30 ± 3.41 Hz; post-injection: 25.09 ± 3.32 Hz, mean ± s.e.m., n=50; p=0.32, Wilcoxon sign-rank test) or control (pre-injection: 26.71 ± 4.89 Hz; post-injection: 31.55 ± 5.27 Hz, mean ± s.e.m., n=31; p=0.20, Wilcoxon sign-rank test) injections into dPL **(Fig. 2B, left**). Also LFP power in V4 was not significantly different between muscimol and control sessions in any of the frequency bands tested (theta (4-7 Hz), p= 0.44; alpha/low-beta (7-15 Hz), p=0.92; beta (15-30 Hz), p= 0.97; low-gamma (30-60 Hz), p=0.82; high-gamma (60-90 Hz), p=0.82; Wilcoxon rank-sum test) **(Fig. 2B, right**). In monkey B, we also recorded spiking activity and LFPs from dPL in addition to LIP and V4 before and after injections with muscimol, or saline, respectively. Baseline firing rates of neurons in dPL were significantly reduced following muscimol injections (pre-injection: 10.69 ± 1.76 Hz; post-injection: 5.48 ± 1.06 Hz, mean ± s.e.m., n=8; p=0.04, Wilcoxon sign-rank test) but not after control injections (pre-injection: 4.33 ± 0.36 Hz; post-injection: 4.46 ± 0.29 Hz, mean ± s.e.m., n=7; p=0.81, Wilcoxon sign-rank test). We also examined high gamma activity (HGA, 60-90Hz), which is often interpreted as a surrogate for mulit-unit activity (Ray, 2008; Ray, 2011; Watson, 2018). Consistent with the reduction in spike rates, the power of HGA in dPL was significantly reduced, following muscimol injections compared to control sessions (p=0.04, Wilcoxon rank-sum test, 8 muscimol sessions, 7 control sessions). No significant changes in LFP power were observed in other frequency bands tested (theta (4-7 Hz), p=0.46, alpha/low-beta (7-15 Hz), p=0.78, beta (15-30 Hz), p=0.96, low gamma (30-60 Hz), p=0.39, Wilcoxon rank-sum test) during muscimol inactivation sessions relative to saline control sessions (**Fig. 2-1**). Taken together, the present results show significant reduction of firing rates and high gamma (60-90Hz) LFP power locally following muscimol injections into dPL, confirming the suggested role of GABAergic neurons in controlling the excitability of local thalamic populations (McCormick, 1987). However, no remote effects on the two interconnected cortical areas were found in terms of baseline firing or LFP power changes as a consequence of dPL inactivation, suggesting that dPL inputs are not critical in maintaining the task independent baseline firing and LFP activity within LIP and V4.

**Figure 2.**
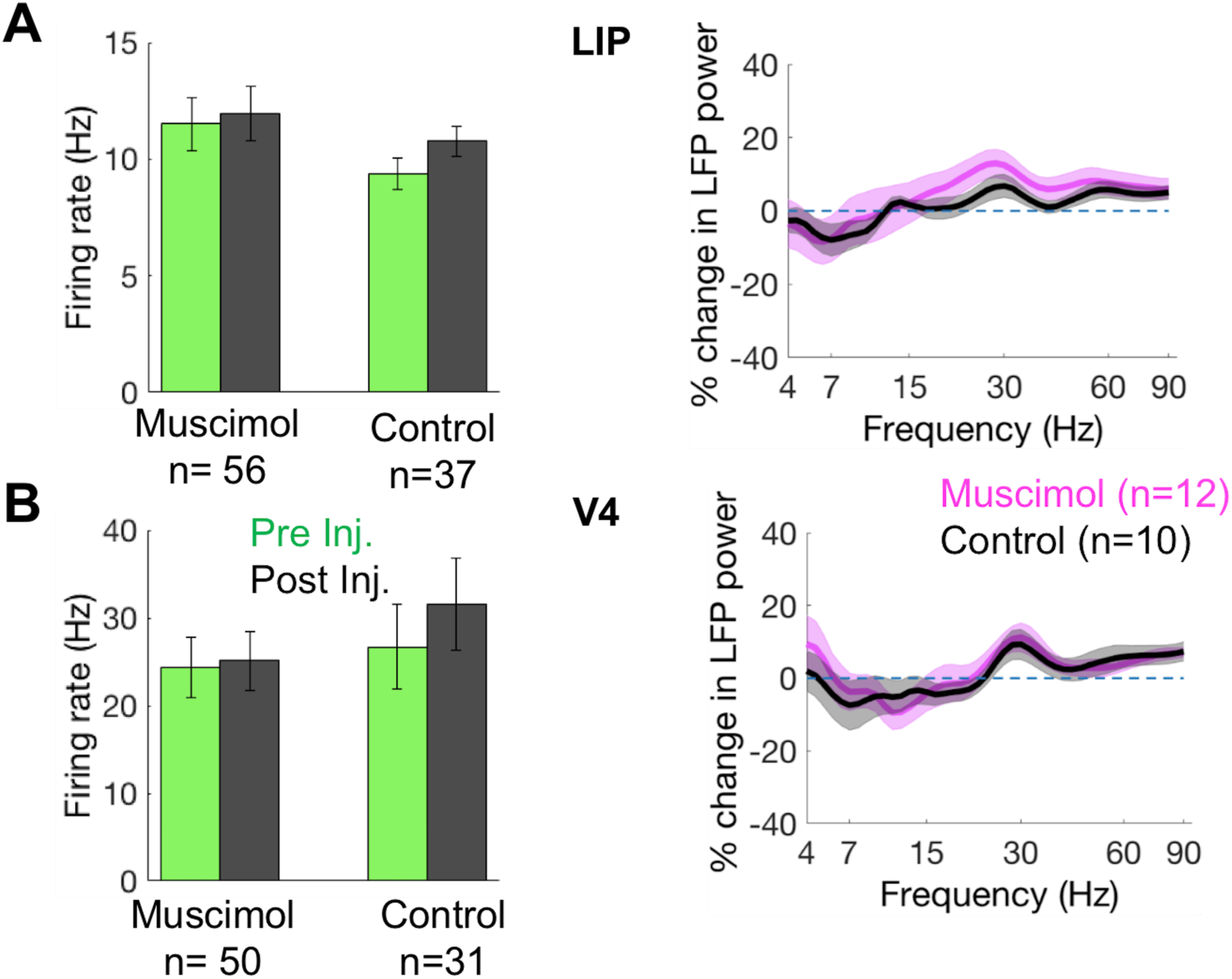
dPL inactivation did not significantly change spike rates and local field potential power in LIP and V4. (A) Left: Changes in LIP spike rates before (green) and after (black) muscimol or control injections into dPL. Right: Percentage change in local field potential (LFP) power in LIP across a frequency range of 4-90 Hz during post vs. pre-injection windows, for muscimol (magenta) and saline control (black) injections. (B) Left: Changes in V4 spike rates before (green) and after (black) muscimol or control injections into dPL. Right: Percentage change in local field potential (LFP) power of V4 across a frequency range of 4-90 Hz during post vs. pre-injection windows, for muscimol (magenta) and saline control (black) injections. Combined data from both monkeys. (Error bars/shaded areas: s.e.m.).

### Effects of pulvinar inactivation on cortico-cortical connectivity

Next we explored the functional role of pulvinar on cortico-cortical connectivity by measuring the phase locking value (PLV) between cortical areas LIP and V4, pre and post muscimol or control injections into dPL. PLV is a measure of LFP phase coherence between areas and is thought to be an effective index for quantifying interareal synchronization, or functional connectivity (Lachaux, 1999). During inactivation sessions, the LFP-LFP phase coherence between LIP and V4 was significantly reduced in theta (4-7 Hz) and alpha/low-beta (7-15 Hz) frequency bands following muscimol injections compared to pre-injection coherence values (theta: p=0.04, alpha/low-beta: p=0.002, n=12 sessions, Wilcoxon sign-rank test). The LFP-LFP coherence was not significantly different during the muscimol sessions in any other frequency bands tested (beta (15-30Hz): p=0.08 and gamma (30-60 Hz): p=0.13, n=12 sessions, Wilcoxon sign-rank test) (**Fig. 3, left**). The frequency band-specific decrease in coherence values was present in both animals (monkey B: alpha/low-beta: p=0.03, p>0.05 in all other frequency bands tested, n=8 sessions, Wilcoxon sign-rank test, monkey R: theta: p=0.04, alpha/low-beta: p=0.05, p > 0.05 in all other frequency bands tested, n=4 sessions, Wilcoxon sign-rank test, one sided). In control sessions, the LIP-V4 phase coherence was not significantly different between pre and post control injection sessions in any of the frequency bands tested in combined data from both animals (theta: p=0.73; alpha: p=0.82; beta: p=0.43; gamma: p=0.25, n=10 sessions, Wilcoxon sign-rank test) (**Fig. 3, right**), or in individual animals (monkey B: p > 0.05 in all frequency bands tested, n=7 sessions, monkey R: p > 0.10, in all frequency bands tested, n=3 sessions, Wilcoxon sign-rank test). The frequency specific LIP-V4 coherence decrease following the muscimol injection into dPL was persistent within the time windows used for analysis (**Fig. 3-1**). Previous studies have suggested, based on correlational analyses, that the pulvinar regulates information processing between interconnected cortical areas depending on task demands in a spatial attention task by modulating functional connectivity in alpha-low beta (8-15 Hz) frequency bands (Saalmann, 2012; Fiebelkorn, 2019a). However, causal evidence for such a role of pulvinar in cortico-cortical synchronization has been lacking. Here, we show that the pulvinar appears to regulate functional connectivity across interconnected areas in low frequency (4-15 Hz) bands, even in the absence of active task conditions.

**Figure 3.**
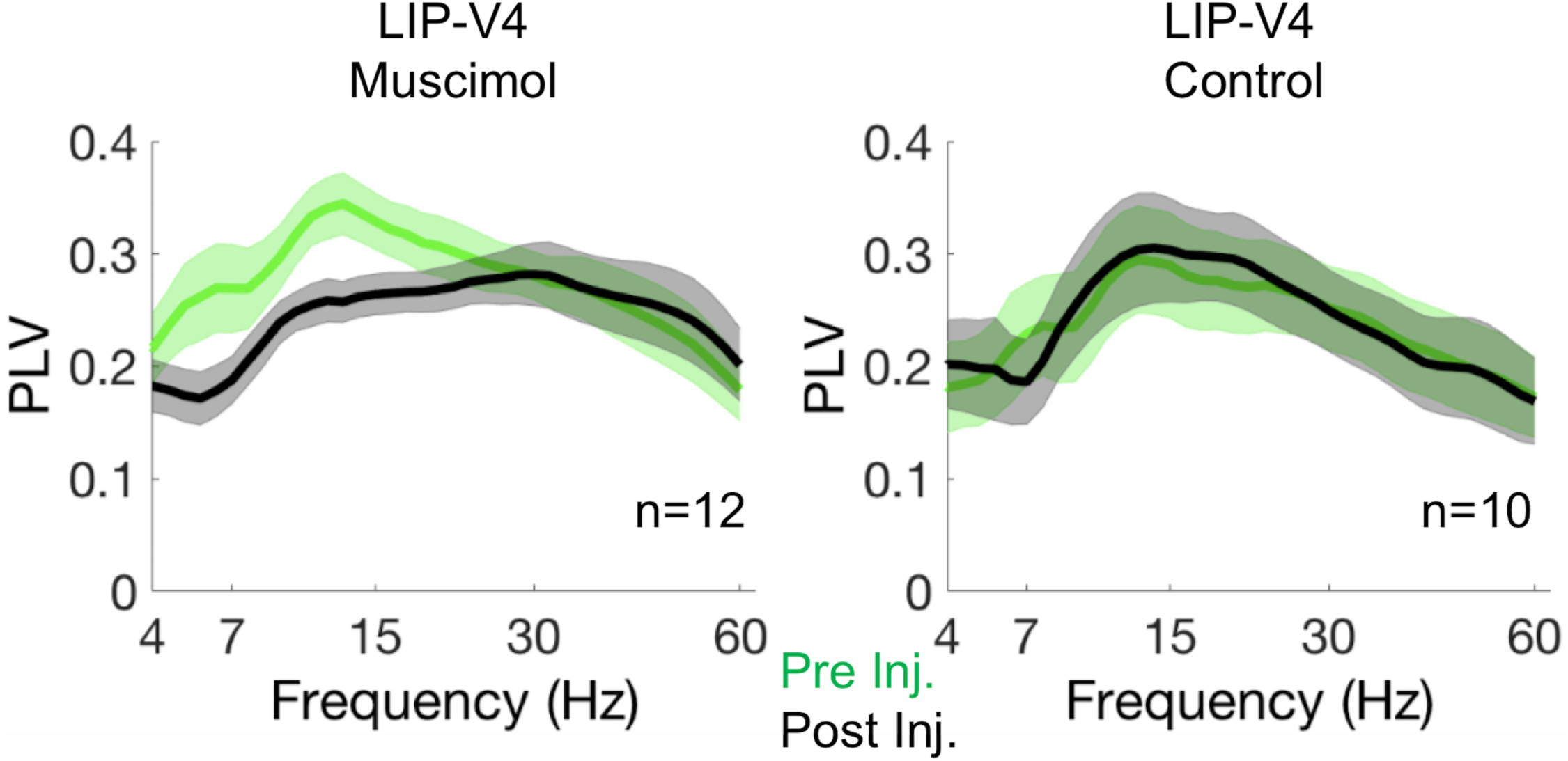
LFP-LFP coherence between LIP and V4 significantly decreased following pulvinar inactivation. Strength of LFP-LFP phase coherence (measured as phase locking value – PLV) between LIP and V4 across a 4-60 Hz frequency range during pre- (green) and post- (black) injection windows, shown separately for muscimol (left) and control (right) sessions. The post-injection LFP-LFP phase coherence between LIP and V4 was significantly lower compared to the pre-injection coherence in theta (4-7 Hz) and alpha/low beta (7-15 Hz) frequency ranges for muscimol, but not for control sessions. Combined data from both monkeys. Shaded areas: s.e.m..

### Changes in pulvino-cortical connectivity following pulvinar inactivation

Thus far, our results show that inactivation of pulvinar caused a significant reduction in the cortico-cortical interactions between LIP and V4 even in the absence of an active task state, while not significantly changing the neuronal firing or LFP power in LIP and V4. In monkey B, we tested the effect of pulvinar inactivation on the phase coherence between pulvinar and cortical areas. Both the pulvinar-LIP and pulvinar-V4 phase coherence values were marginally reduced in alpha/low-beta (7-15 Hz) frequency bands following the muscimol injections into dPL (pulvinar-LIP, p=0.05, (**Fig. 3-2 A, left**); pulvinar-V4, p=0.06, (**Fig. 3-2 B, left**), p > 0.05 in all other frequency bands tested, n=8 sessions, Wilcoxon sign-rank test), but not after the control injections (p > 0.05, in all frequency bands tested, both for pulvinar-LIP and pulvinar-V4 coherences, n=7 sessions, Wilcoxon sign-rank test) (**Fig. 3-2 A, B, right**). These results confirm that the pulvino-cortical connectivity that is mediated through neuronal synchronization in the alpha/low beta frequencies observed during attention tasks is present even in the absence of an active task state. However, the LFP-LFP phase synchronization is not informative regarding the direction of the observed inter-areal interaction. Therefore, we measured spike field coherence (SFC) between pulvinar and cortical areas to further explore pulvino-cortical interactions. SFC is an indirect measure of the relationship between spiking outputs from one region to the synaptic inputs (LFP) of a second region, and thus provides a useful tool for exploring inter-areal input-output interactions (Fries, 2001; Pesaran, 2010). A frequency-specific increase in SFC can be interpreted as arising from the area that produces spiking output and being directed at LFP signals of the input area (Pesaran, 2010). We measured SFC between pulvinar and cortical areas in both directions (from pulvinar to cortex and from cortex to pulvinar), for pre- and post-injection windows. SFC between pulvinar spiking activity and LIP LFP phase post- vs. pre-injection was significantly lower after muscimol injection in theta-alpha/low-beta (4-15 Hz) frequencies, as compared to the corresponding difference after control injections (p=0.01, p > 0.05 in beta and gamma frequency ranges, Wilcoxon rank-sum test) (**Fig. 4A, left**). Similarly, the difference in SFC between pulvinar spikes and V4 LFP phase was significantly reduced in theta-alpha/low-beta frequencies after dPL inactivation compared to corresponding measures after control sessions (p=0.03, p > 0.05 in beta and gamma frequency ranges, Wilcoxon rank-sum test) (**Fig. 4B, left**). Interestingly, the difference in SFC between LIP spikes and pulvinar LFP phase significantly increased in a similar theta-alpha/low-beta frequency range following the muscimol injections into dPL, as compared to control injections into dPL (p=0.02, p > 0.05 in beta and gamma frequency ranges, Wilcoxon rank-sum test) (**Fig. 4A, right**), indicating an increase in spike-field coherence between LIP spikes and pulvinar phase following the muscimol injections into dPL. No such increase in SFC was observed between V4 spikes and pulvinar LFP phase in this or any other frequencies (theta-alpha/low-beta: p=0.69; other frequencies: p > 0.05; Wilcoxon rank-sum test) (**Fig. 4B, right**). Since the spike field measures provide indirect evidence for the directionality of inter-areal influences (i.e. spikes as output signals from an area and LFPs input signals into an area), the current SFC results suggest a significant decrease in network connectivity from pulvinar to cortical areas (LIP and V4) in low frequency range (4-15Hz), following the muscimol inactivation of dPL. Our results also suggest that the network connectivity from LIP to pulvinar, but not from V4 to pulvinar, is significantly increased following dPL inactivation. Taken together, these results are consistent with and complement our main finding that pulvinar inactivation interferes with cortico-cortical interactions mediated in low frequencies (4-15Hz).

**Figure 4.**
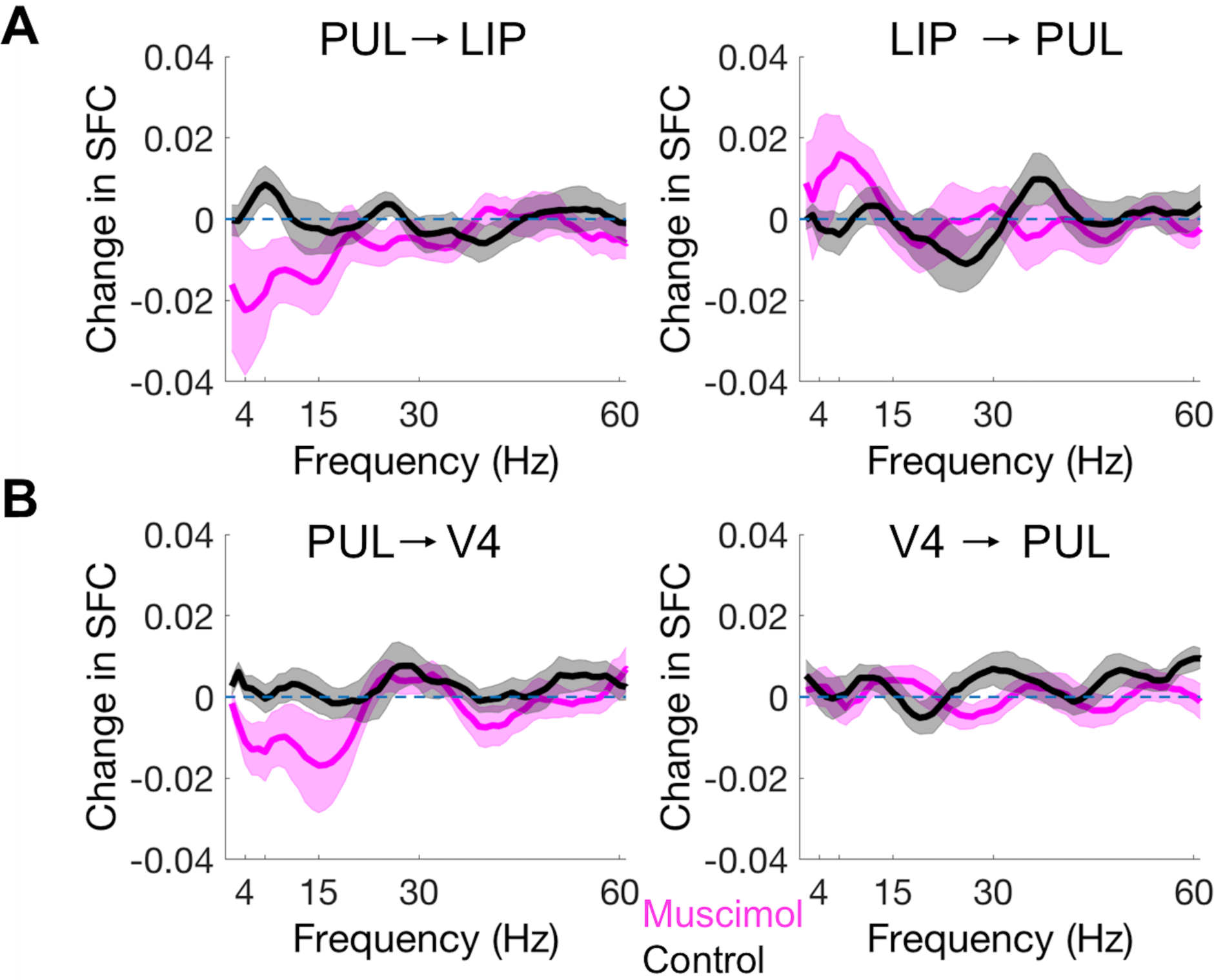
Changes in spike-field coherence (SFC) between pulvinar and cortical areas following dPL injections. (A) Left: Change in SFC between pulvinar spikes and LIP LFP phase over a frequency range of 4-60 Hz during post vs. pre-injection windows for muscimol (magenta) and control (black) sessions. The dPL spike-LIP phase coherence decreased significantly in the 4-15 Hz range following muscimol injections into dPL. Right: Change in SFC between LIP spikes and pulvinar LFP phase during post vs. pre-injection windows for muscimol (magenta) and control (black) sessions. The LIP spike-pulvinar phase coherence increased significantly in the 4-15 Hz range following muscimol injections. (B) Left: change in SFC between pulvinar spikes and V4 LFP phase during post vs. pre-injection windows for muscimol (magenta) and control (black) sessions. The dPL spike-V4 phase coherence decreased significantly in the 4-15 Hz range following muscimol injections into dPL. Right: Change in SFC between V4 spike and pulvinar LFP phase during post vs. pre-injection windows for muscimol (magenta) and control (black) sessions. Shaded areas: s.e.m..

## Discussion

We performed simultaneous recordings from dorsal (LIP) and ventral (V4) areas of the visual system, while reversibly inactivating the anatomically interconnected subdivision of the pulvinar (dPL) in monkeys during passive viewing conditions. The results show a significant reduction in local field phase coherence between LIP and V4 in low frequencies (4-15 Hz) following muscimol injection into dPL. This finding provides causal evidence for a role of pulvinar in regulating cortico-cortical connectivity even in the absence of an active task state. Our results suggest that the pulvinar’s influence on cortico-cortical interactions is not limited to specific task requirements, but is part of more basic functional connectivity that can be modulated by task demands. At the local level, no significant changes in firing rates or LFP power were observed in LIP or in V4 following dPL inactivation, indicating that dPL inputs are not critical in maintaining local passive response properties within extrastriate, or parietal visual areas. However, significant decreases in synchronization between pulvinar spikes and cortical LFP phase were observed both in LIP and V4, following dPL inactivation. The reduction in pulvino-cortical spike field coherence was specific for theta-alpha/low-beta frequencies (4-15Hz), suggesting a role for pulvinar spike inputs in frequency-specific synchronization of cortico-cortical interactions.

Electrophysiological studies on the pulvinar have provided evidence for a role in regulating information transmission between ventral visual areas (Saalmann, 2012), or between higher-order cortical areas (e.g. FEF and LIP) (Fiebelkorn, 2019a) during spatial attention tasks. These studies explored the role of pulvinar under specific task conditions, leaving open the question as to whether the pulvinar influence on cortico-cortical functional connectivity is related to engagement in specific tasks, or rather a more basic property of large-scale network connectivity. Studies in humans have suggested functional connectivity between pulvinar and cortical areas even in the absence of any active task structure, based on BOLD responses (Stein, 2000; Barron, 2015; Terpou, 2018). Watching short movie clips during the data collection keeps subjects more engaged and has been shown to be a reliable method for obtaining functional connectivity data both in monkeys (Mantini, 2013) and in humans (Vanderwal, 2015; Vanderwal, 2019). In the current study, we adopted this method and recordings were performed while animals freely viewed short movie clips.

Previous immunohistochemistry studies have shown the distribution of GABAergic neurons across different mammalian thalamic structures (Sivilotti, 1991; Arcelli, 1997). For the pulvinar, moderate to high densities of GABAergic neurons were observed in lateral, medial and inferior subdivisions of pulvinar in monkeys (Hunt, 1991; Huntsman, 1996). These studies have suggested a crucial role for GABAergic neurons in regulating thalamocortical circuits by changing the excitability of local thalamic populations (McCormik, 1987; Tremblay, 2016). Muscimol, a GABA_A_ agonist which causes local hyperpolarization, has been extensively used to reversibly inactivate brain regions and to explore causal relationships (Petersen, 1987; Desimone, 1990; Wilke, 2010). Previous inactivation studies probing a causal role for the pulvinar in influencing local cortical responses have shown various results on baseline, visually-evoked, or task-related responses. In anesthetized preparations, both baseline firing rates and visually-evoked responses were found to be significantly reduced in V1 following muscimol injections into the lateral pulvinar (Purushothaman, 2012). In contrast to V1 responses in anesthetized recordings, baseline firing rates in V4 neurons were marginally increased in awake monkeys performing an attention task, following muscimol injections into the ventro-lateral pulvinar (Zhou, 2016). The visually-evoked responses, however, were significantly decreased following ventro-lateral pulvinar inactivation (Zhou, 2016). In the current study, we did not find any significant changes in task independent baseline firing rates or LFP power following muscimol injection into dPL. It is possible that the differences in preparation (anesthetized vs. awake behaving) account for the differences in pulvinar inactivation effects on baseline firing rates between V1 and V4, as observed in these studies. However, alternatively, it is possible that there is a fundamental difference between pulvino-cortical loops involving primary visual versus extrastriate cortex, as suggested by studies in rodent somatosensory cortex (Sherman, 2007; Theyel, 2009). This possibility needs to be probed in future studies in primates. Taken together, the effects of pulvinar inactivation on baseline firing rates, observed previously (Zhou, 2016) and in the present study suggest that pulvinar inputs do not appear to impact local neuronal response properties in the absence of active task demands in extrastriate area V4 and in parietal area LIP.

Previous studies suggest that pulvinar coordinates information transmission across cortical areas by synchronizing their activity in alpha/low beta frequencies (Saalmann, 2012). For example, during a spatial attention task, the LFP-LFP coherence between V4 and TEO, as well as V4 and pulvinar and TEO and pulvinar was significantly increased when attention was deployed at a RF as compared to away from it, predominantly in alpha/low-beta (8-15 Hz) frequencies. Granger analysis suggested a causal influence from pulvinar to cortex (Saalmann, 2012). Here, we provide critical empirical evidence for this notion by perturbing the V4-LIP-dPL network with an actual causal manipulation and by demonstrating a decrease in neural synchronization between V4 and LIP in a similar frequency band following muscimol inactivation of the interconnected dPL. Thus, perturbation of pulvinar appears to affect primarily the functional connectivity of interconnected network nodes, substantiating the notion that pulvinar plays a causal role in coordinating inter-areal functional interactions in cortex.

Recent studies performing simultaneous recordings from pulvinar and cortical nodes of the fronto-parietal attention network (FEF and LIP) have further established a role of pulvinar in regulating cortico-cortical interactions in spatial attention tasks (Fiebelkorn, 2019a). It was shown that the medial pulvinar rhythmically engaged and disengaged with the cortical nodes (FEF and LIP) during the allocation of attention in alpha/low beta (14-24 Hz) frequencies. During periods of attentional engagement, Granger analyses suggested that information flowed from pulvinar to cortex via the transthalamic pathway, thereby facilitating visual processing. The direction reversed from cortex to pulvinar during periods of attentional disengagement, suggesting a shutdown of the transthalamic pathway (Fiebelkorn, 2019a, b). In the current study, we found a significant decrease in spike-field coherence between pulvinar spikes and LFP phases in V4 and LIP following muscimol inactivation specifically in low frequency (4-15 Hz) ranges. Interestingly, the spike-field coherence between LIP spikes and pulvinar phase was found to be increased following pulvinar inactivation in similar frequency ranges. With the general notion of spikes being output signals and LFPs being input signals (Pesaran, 2010), the current SFC findings suggest a reversal of information flow from cortex to pulvinar following the pulvinar inactivation. Thus, the influence of higher-order cortex on pulvinar appeared to dominate the functional interactions between pulvinar and LIP, similar to what has been observed during states of disengagement in these recent studies (Fiebelkorn, 2019a, b), suggesting that the rhythmic reweighting of functional interactions between thalamus and cortex is a more general property of the visual large-scale network and may be modulated rather than generated during cognitive task states.

In summary, our results reveal a causal role of pulvinar in regulating cortico-cortical interactions even in the absence of an active task structure. The results suggest that pulvinar inputs may not impact baseline responses, at least in extrastriate cortex, but rather are critical in synchronizing responses between different interconnected cortical nodes of a cortical large-scale network. Future studies of pulvinar inactivation and simultaneous recordings from nodes of the fronto-parietal network under behavioral tasks will be required to further establish causal mechanisms of task-dependent selective cortical engagement by pulvinar during attention and other cognitive tasks.

## Acknowledgments

This work was supported by grants from NEI (2R01EY017699) and NIMH (2R01MH064043, P50MH109429).

---

**Figure 1-1.**
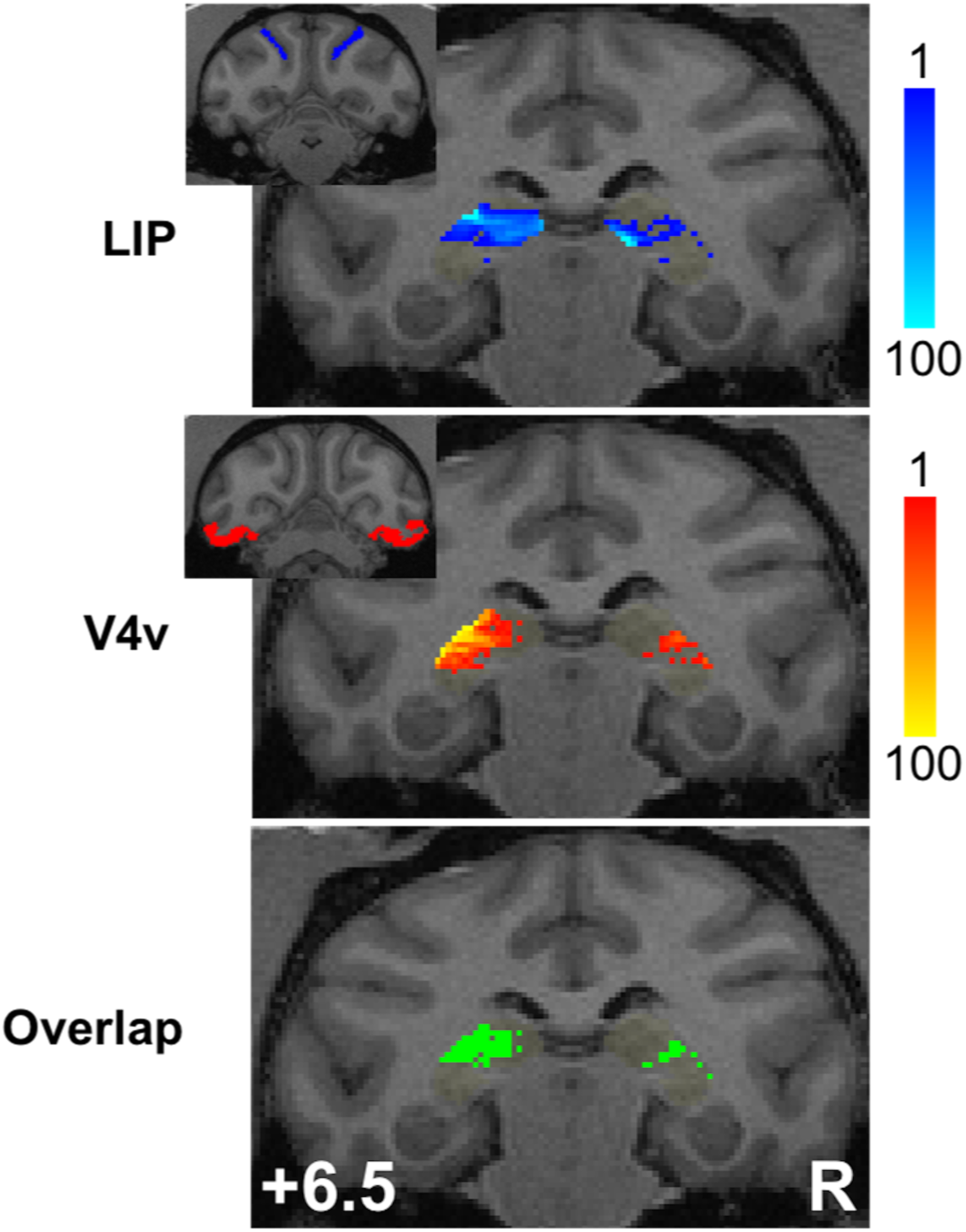
Zone of convergent connectivity within lateral pulvinar. Probabilistic tractography was performed in monkey B to determine the convergent zone for likely fiber pathways from LIP and V4 (see inlets in blue and red, respectively) to pulvinar. Regions color-coded in green indicate convergent projection zones of LIP and V4 within the lateral pulvinar.

**Figure 1-2.**
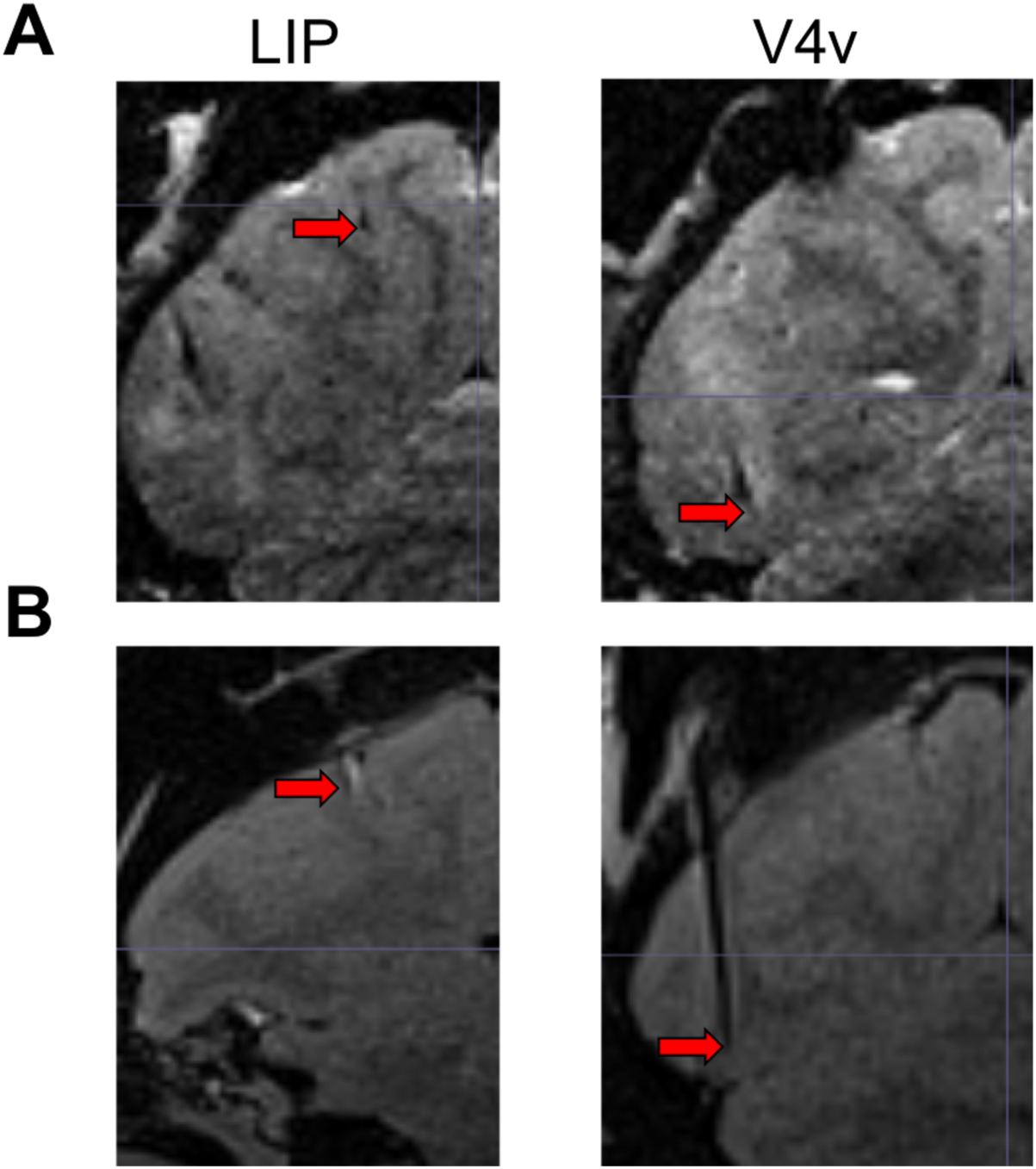
Localization of electrophysiological recording locations for LIP and V4. A) T1 MRI images showing the electrode positions for LIP (left) and V4 (right) for Monkey B B) T1 MRI images showing the electrode positions for LIP (left) and V4 (right) for Monkey R. The red arrows indicate the starting tip position of the tungsten electrodes placed for position confirmation. The LIP location ranged from AP -2 to AP -5 while V4 locations ranged from AP -3 to AP -6, distances in mm caudal to ear bar zero (EBZ).

**Figure 2-1.**
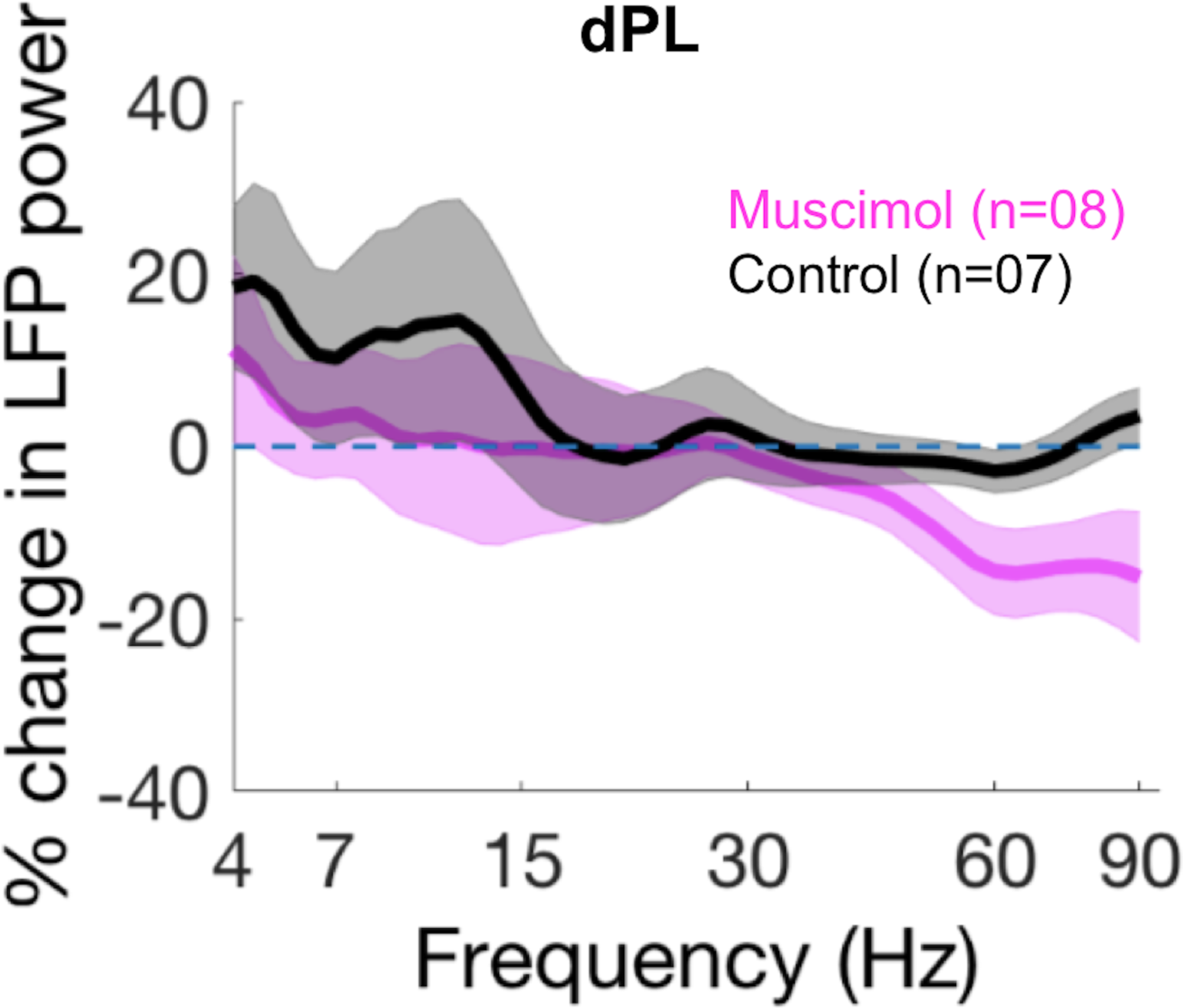
dPL inactivation significantly decreased high gamma local field potential power within pulvinar. Percentage change in local field potential (LFP) power in dPL during post-injection period relative to pre-injection period, across a frequency range of 4-90 Hz, for muscimol (magenta) and saline control (black) injections. The task independent baseline LFP power in dPL significantly decreased in the gamma frequency range (60-90 Hz), but not in other frequency ranges, following muscimol injections, compared to saline control injections. Shaded areas indicate s.e.m.

**Figure 3-1.**
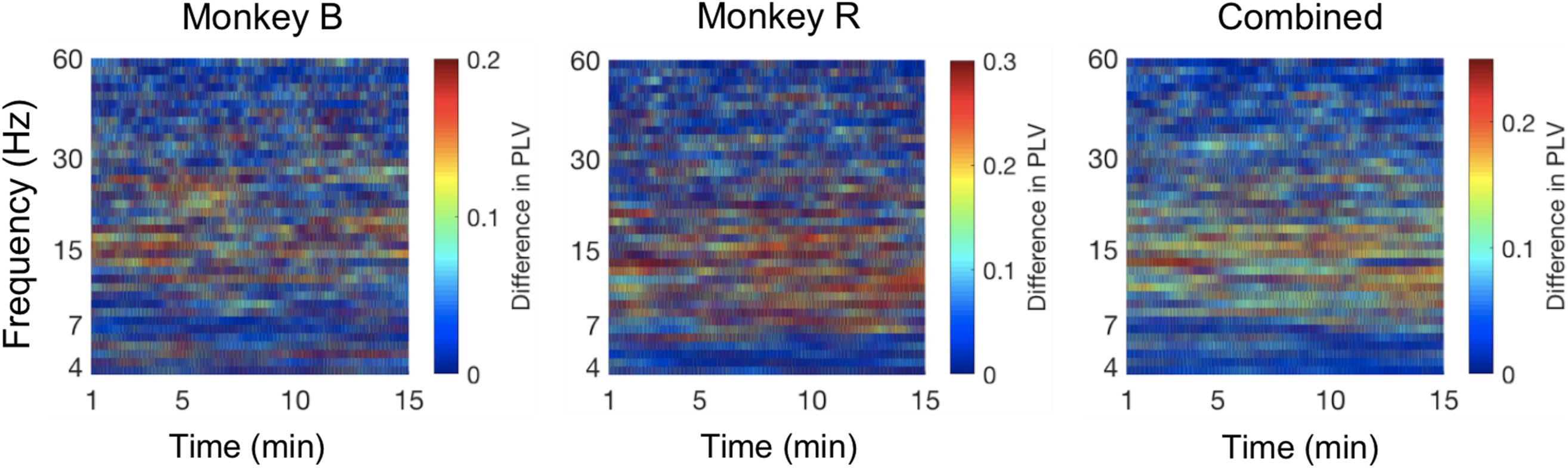
Time resolved decrease in LFP-LFP phase coherence between LIP and V4 following dPL inactivation in individual animals. The difference in pre- and post-injection window LFP-LFP phase coherence between LIP and V4 across a 4-60 Hz frequency range plotted across the window period (15min) for monkey B, monkey R and combined between monkeys.

**Figure 3-2.**
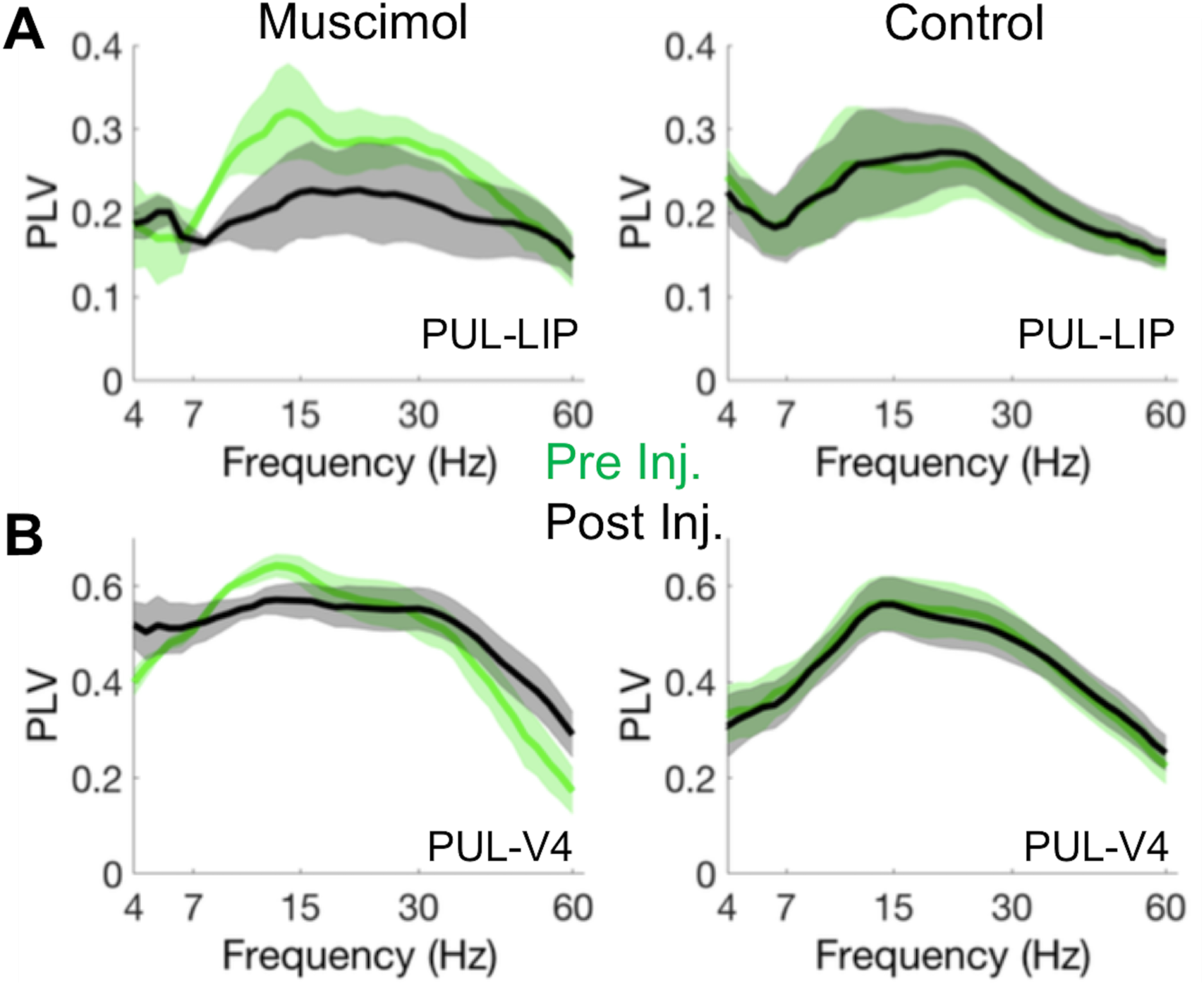
Decrease in LFP-LFP phase coherence between pulvinar and cortical areas following dPL inactivation. A) LFP-LFP phase coherence between pulvinar and LIP across a 4-60 Hz frequency range during pre- (green) and post- (black) injection windows for muscimol (left) and control (right) sessions. B) PLV between pulvinar and V4 across a 4-60 Hz frequency range, for pre- (green) and post- (black) injection windows for muscimol (left) and control (right) sessions. Shaded areas: s.e.m.

## Notes

**Conflict of interest statement** The authors declare no conflicts of interests

